# The eardrums move when the eyes move: A multisensory effect on the mechanics of hearing

**DOI:** 10.1101/156570

**Authors:** K. G. Gruters, D. L. K. Murphy, Cole D. Jenson, D. W. Smith, C. A. Shera, J. M. Groh

## Abstract

Interactions between sensory pathways such as the visual and auditory systems are known to occur in the brain, but where they *first* occur is uncertain. Here we show a novel multimodal interaction evident at the eardrum. Ear canal microphone measurements in humans (*n*=19 ears in 16 subjects) and monkeys (*n*=5 ears in 3 subjects) performing a saccadic eye movement task to visual targets indicated that the eardrum moves in conjunction with the eye movement. The eardrum motion was oscillatory and began as early as 10 ms before saccade onset in humans or with saccade onset in monkeys. These eardrum movements, which we dub Eye Movement Related Eardrum Oscillations (EMREOs), occurred in the absence of a sound stimulus. The EMREOs’ amplitude and phase depended on the direction and horizontal amplitude of the saccade. They lasted throughout the saccade and well into subsequent periods of steady fixation. We discuss the possibility that the mechanisms underlying EMREOs create eye movement-related binaural cues that may aid the brain in evaluating the relationship between visual and auditory stimulus locations as the eyes move.

**SIGNIFICANCE STATEMENT:** The peripheral hearing system contains several motor mechanisms that allow the brain to modify the auditory transduction process. Movements or tensioning of either the middle-ear muscles or the outer hair cells modify eardrum motion, producing sounds that can be detected by a microphone placed in the ear canal (e.g. as otoacoustic emissions). Here, we report a novel form of eardrum motion produced by the brain via these systems -- oscillations synchronized with and covarying with the direction and amplitude of saccades. These observations suggest that a vision-related process modulates the first stage of hearing. In particular, these eye-movement related eardrum oscillations may help the brain connect sights and sounds despite changes in the spatial relationship between the eyes and the ears.

## INTRODUCTION

Visual information can aid hearing, such as when lip reading cues facilitate speech comprehension. To derive such benefits, the brain must first link visual and auditory signals that arise from common locations in space. In species with mobile eyes (such as humans and monkeys), visual and auditory spatial cues bear no fixed relationship to one another but change dramatically and frequently as the eyes move—about three times per second over an 80 degree range of space. Accordingly, considerable effort has been devoted to determining where and how the brain incorporates information about eye movements into the visual and auditory processing streams (1). In the primate brain, all of the regions previously evaluated have shown some evidence that eye movements modulate auditory processing (IC: 2, 3-6) (auditory cortex: 7, 8, 9) (parietal cortex: 10, 11, 12) (superior colliculus: 13, 14-17). Such findings raise the question of where in the auditory pathway eye movements *first* impact auditory processing. In this study, we tested whether eye movements affect processing in the auditory periphery.

The auditory periphery possesses at least two means of tailoring its processing in response to descending neural control (Figure 1): (1) The middle ear muscles (MEMs)—the stapedius and tensor tympani—attach to the ossicles that connect the eardrum to the oval window of the cochlea. Contraction of these muscles tugs on the ossicular chain, modulating middle-ear sound transmission and moving the eardrum. (2) Within the cochlea, the outer hair cells (OHCs) are mechanically active and modify the motion both of the basilar membrane, and, through mechanical coupling via the ossicles, the eardrum (i.e., otoacoustic emissions). In short, the actions of the MEMs and OHCs affect not only the response to incoming sound but also transmit vibrations backwards to the eardrum. Both the MEMs and OHCs are subject to descending control by signals from the central nervous system (see refs: 18, 19, 20) for reviews), allowing the brain to adjust the cochlear encoding of sound in response to previous or ongoing sounds in either ear and based on global factors, such as attention (21–27). The collective action of these systems can be measured in real time with a microphone placed in the ear canal (28). We used this technique to study whether the brain sends signals to the auditory periphery concerning eye movements, the critical information needed to reconcile the auditory spatial and visual spatial worlds.

**Figure 1.**
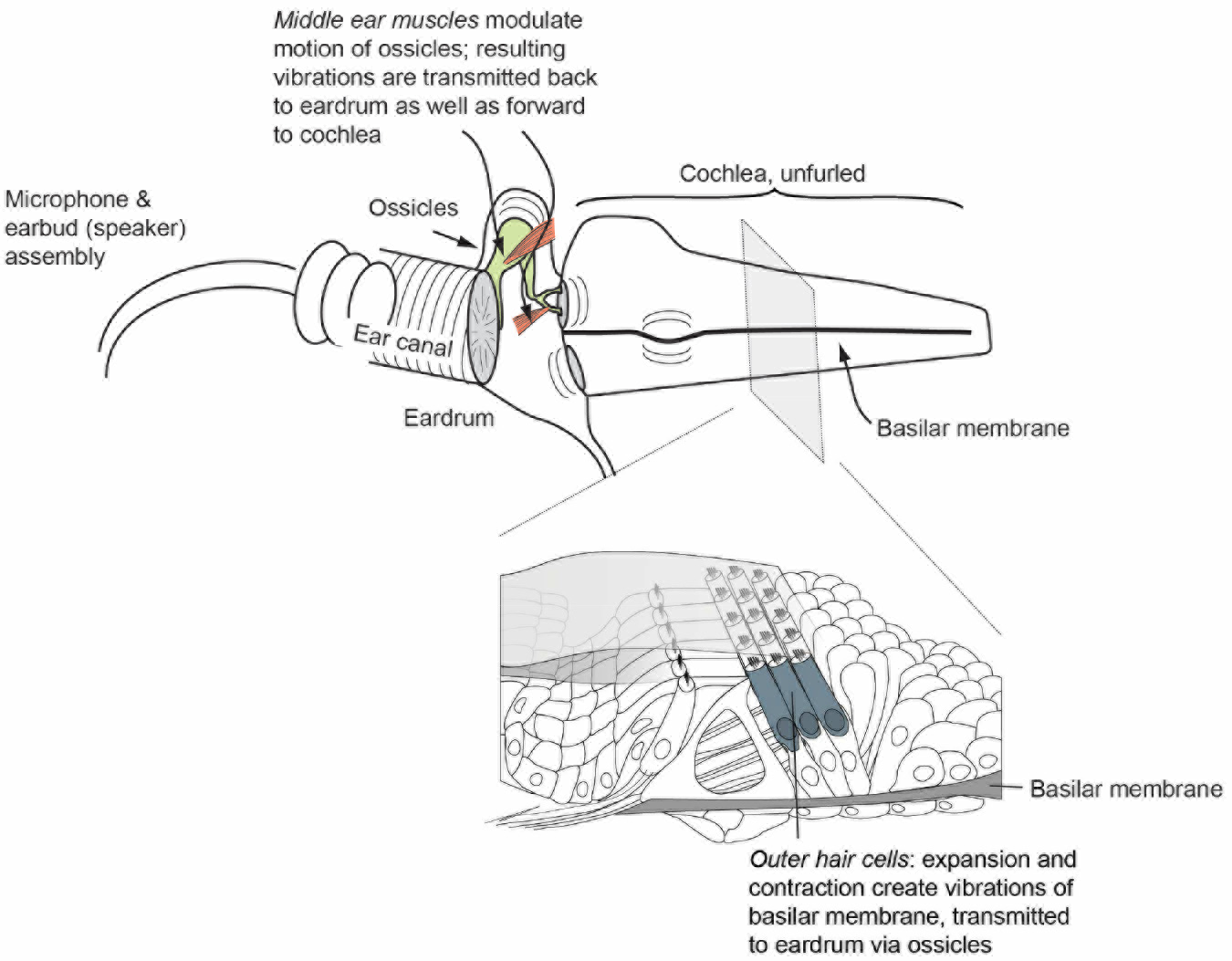
Motile cochlear outer hair cells (OHCs) expand and contract in a way that depends both on the incoming sound and on descending input received from the superior olivary complex in the brain. OHC motion moves the basilar membrane, and subsequently the eardrum via fluid/mechanical coupling of these membranes through the ossicular chain. The middle-ear muscles also pull on the ossicles, directly moving the eardrum. These muscles are innervated by motor neurons near the facial and trigeminal nerve nuclei, which receive input from the superior olive bilaterally. In either case, eardrum motion can be measured with a microphone in the ear canal.

**Figure 2.**
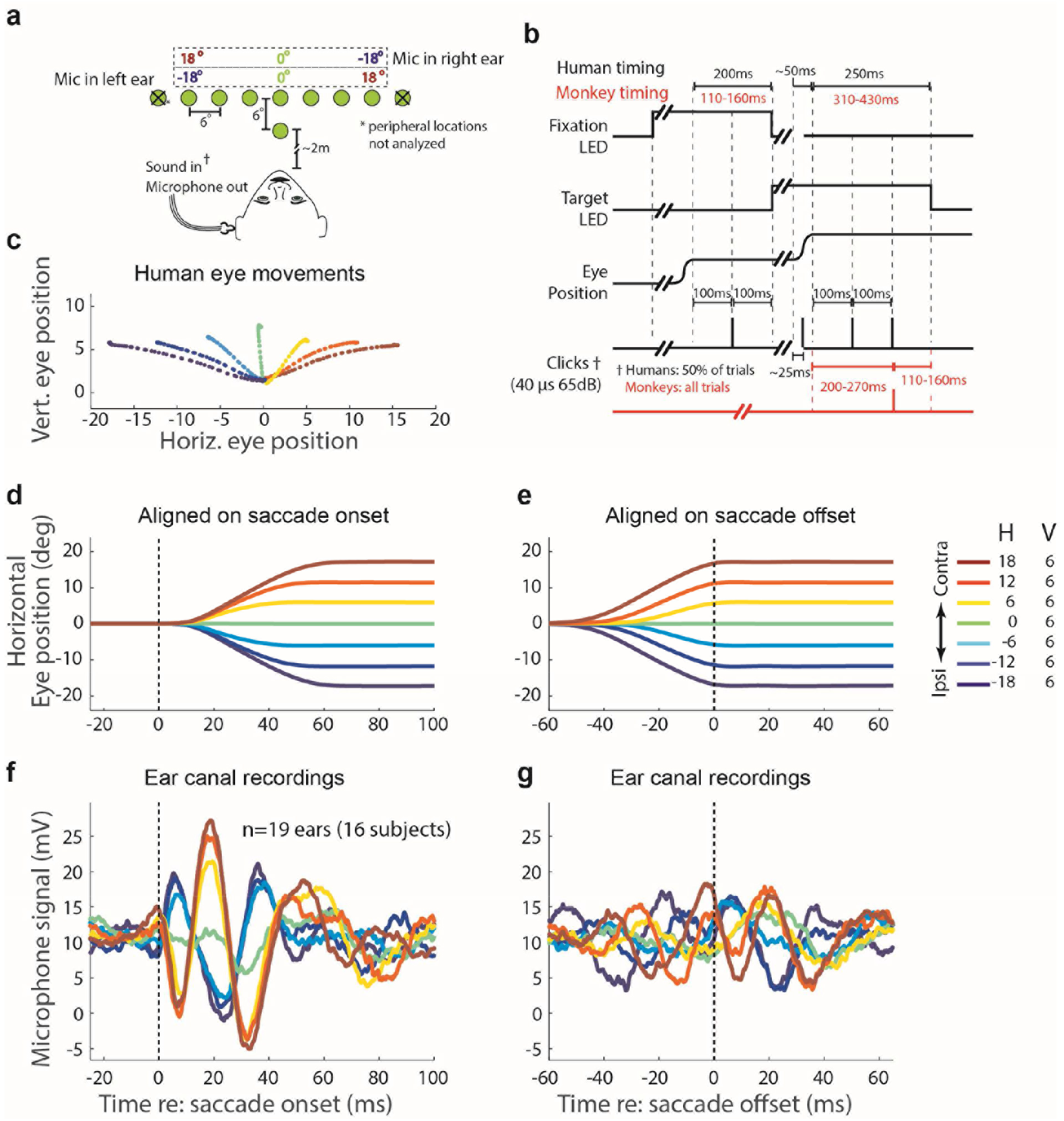
Experimental design and results. **a.** Recordings for all subjects were made via a microphone in the ear canal set into a custom-fit ear bud. On each trial, a subject fixated on a central LED, then made a saccade to a target LED (−24° to +24° horizontally in 6° increments and 6° above the fixation point) without moving his/her head. The ±24° locations were included on only 4.5% of trials and were excluded from analysis (see Methods); other target locations were equally likely (~13% frequency). **b.** Humans (black text) received randomly interleaved silent and click trials (50% each). Clicks were played via a sound transducer coupled with the microphone at four times during these trials: during the initial fixation and saccade, and at 100ms and 200ms after target fixation. Monkeys’ trials had minor timing differences (red text) and all trials had one click at 200-270 ms after target fixation (red click trace). **c.** Average eye trajectories for one human subject and session for each of the included targets, colors indicate saccade target locations from ipsilateral (blue) to contralateral (red). **d,e.** Mean eye position as a function of time, aligned on saccade onset (d) and offset(e). **f,g.** Mean microphone recordings of air pressure in the ear canal, aligned on saccade onset (f) and offset (g) indicate that the eardrum oscillates in conjunction with eye movements. The phase and amplitude of the oscillation varied with saccade direction and amplitude respectively. The oscillations were on average larger when aligned on saccade onset than when aligned on saccade offset.

## >RESULTS

### The eardrum moves with saccades

Sixteen humans executed saccades to visual targets varying in horizontal position (Figure 2A). Half of the trials were silent whereas the other half incorporated a series of task-irrelevant clicks presented before, during, and at two time points after the saccade (Figure 2B). This allowed us to compare the effects of eye movement-related neural signals on the auditory periphery in silence as well as in the presence of sounds often used to elicit otoacoustic emissions.

We found that the eardrum moved when the eyes moved, even in the absence of any externally delivered sounds. Figure 2C-D shows the average eye position as a function of time for each target location for the human subjects on trials with no sound stimulus, aligned on saccade onset (Figure 2C) and offset (Figure 2D). The corresponding average microphone voltages are similarly aligned and color-coded for saccade direction and amplitude (Figure 2E, F). The microphone readings oscillated time-locked to both saccade onset and offset with a phase that depended on saccade direction. When the eyes moved toward a visual target contralateral to the ear being recorded, the microphone voltage deflected positively, indicating a change in ear canal pressure, beginning about 10 ms prior to eye movement onset. This was followed by a more substantial negative deflection, at about 5 ms after the onset of the eye movement, after which additional oscillatory cycles occurred. The period of the oscillation is typically about 30 ms (~33Hz). For saccades in the opposite (ipsilateral) direction, the microphone signal followed a similar pattern but in the opposite direction: the initial deflection of the oscillation was negative. The amplitude of the oscillations varied with the amplitude of the saccades, with larger saccades associated with larger peaks than those occurring for smaller saccades.

Comparison of the traces aligned on saccade onset vs saccade offset reveals that the oscillations continue into at least the initial portion of the period of steady fixation that followed saccade onset. The EMREO observed following saccade offset was similar in form to that observed during the saccade itself when the microphone traces were aligned on saccade onset, maintaining their phase and magnitude dependence on saccade direction and length. The fact that this post-saccadic continuation of the EMREO is not seen when the traces are aligned on saccade onset suggests a systematic relationship between the offset of the movement and the phase of the EMREO, such that variation in saccade duration obscures the ongoing nature of the EMREO.

**Figure 3.**
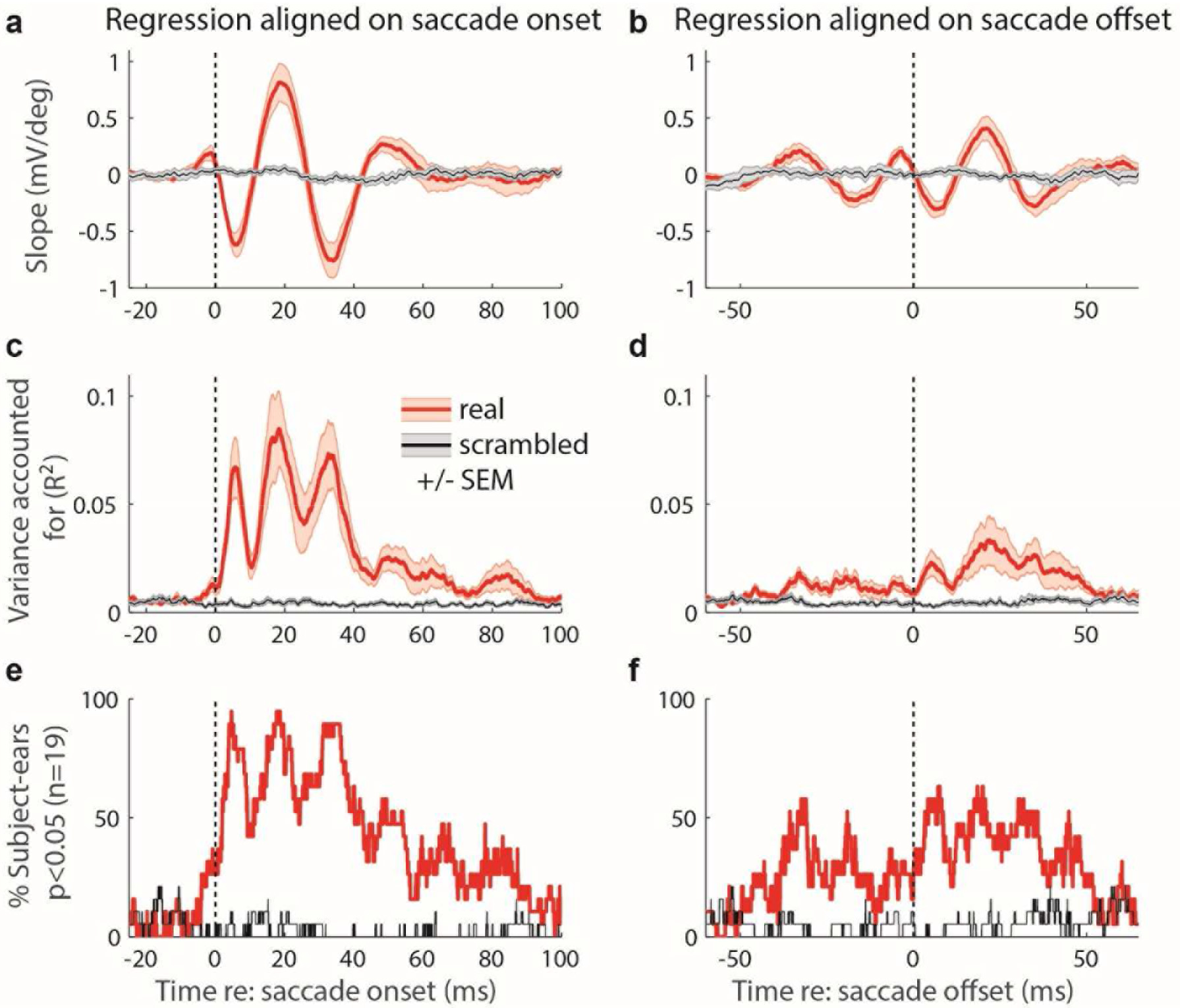
Regression results for data aligned to saccade onset (a, c, e) and offset (b, d, f). **a, b.** Mean ±S.E.M. slope of regression of microphone voltage vs. saccade target location (conducted separately for each subject then averaged across the group) at each time point for real (red) vs. scrambled (gray) data. In the scrambled Monte Carlo analysis, the true saccade targets locations were shuffled and arbitrarily assigned to the microphone traces for individual trials. **c, d.** Proportion of variance accounted for by regression fit (R^2^). **e, f.** Percent of subject-ears showing p <0.05 for the corresponding time point. See Methods for additional details.

To obtain a portrait of the statistical significance of the relationship between the direction and amplitude of the eye movements and the observed microphone measurements of ear-canal pressure across time, we calculated a regression of microphone voltage vs. saccade target location for each 0.04 ms sample from 25 ms before to 100 ms after saccade onset. The regression was conducted separately for each individual subject. Since this involved many repeated statistical tests, we compared the real results with a Monte Carlo simulation in which we ran the same analysis but scrambled the relationship between each trial and its true saccade target location (see Supplemental Methods section for details). As shown in Figure 3A, the slope of the regression involving the real data (red trace, averaged across subjects) frequently deviates from zero, indicating a relationship between saccade amplitude and direction and microphone voltage. The value of the slope oscillates during the saccade period, beginning 9 ms prior to and continuing until about 60 ms after saccade onset, matching the oscillations evident in the data in Figure 2E. In contrast, the scrambled data trace (gray) is flat during this period. Similarly, the effect size (*R*^2^ values) of the real data deviate from the scrambled baseline in a similar but slightly longer timeframe, dropping back to the scrambled data baseline at 75-100 ms after saccade onset (Figure 3C). Figure 3E shows the percent of subjects showing a statistically significant (*p*<0.05) effect of saccade target location at each time point. This curve reaches 80-100% during the peaks of the oscillations observed at the population level. A similar, though weaker, dependence of the EMREO on target location was observed when the recordings were synchronized to saccade offset (Figure 3B,D, F). We repeated this test in an additional dataset (human data set II, see methods) involving finer grained sampling within a hemifield (i.e. analyzing target locations within the contralateral or ipsilateral hemifield). This analysis confirmed the relationship between the microphone voltage and saccade amplitude, as well as saccade direction (Supplementary Figure 1).

### EMREO phase is related to relative, not absolute, saccade direction

Comparison of the EMREOs in subjects who had both ears recorded confirmed that the direction of the eye movement relative to the recorded ear determines the phase pattern of the EMREO. Figure 4 shows the results for one such subject. When the left ear was recorded, leftward eye movements were associated with a positive voltage peak at saccade onset (Figure 4A, blue traces), whereas when the right ear was recorded, that same pattern occurred for rightward eye movements (Figure 4B, blue traces). Translating this into eardrum motion, the implication is that for a given moment during a given saccade, the eardrums bulge inward in one ear while bulging outward in the other ear. Which ear does what when is determined by the direction of the eye movement relative to the recorded ear (contralaterality/ipsilaterality), not whether it is directed to the left vs right in space. The other subjects tested with both ears showed similar patterns, and those tested with one ear were also generally consistent (although there were some individual idiosyncrasies in timing and amplitude; see Supplementary Figure 2 for the remaining individual subject data).

**Figure 4.**
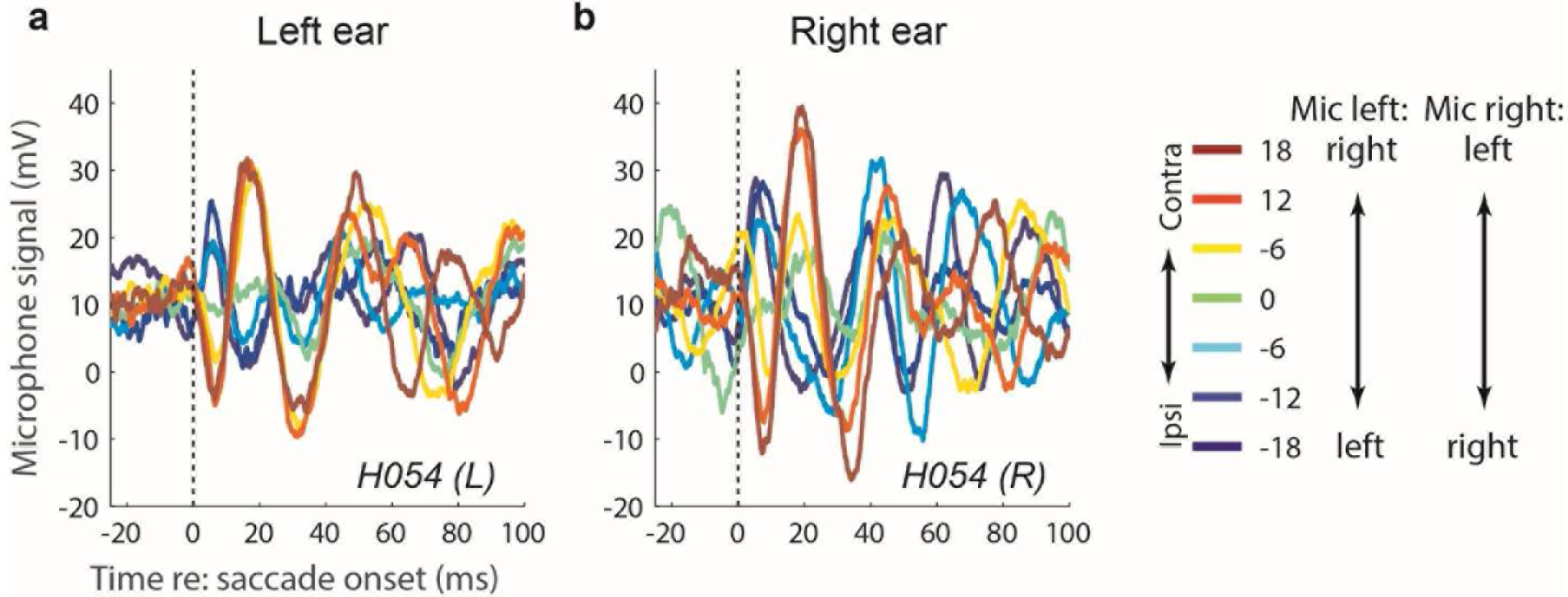
Recordings in left and right ears in an individual human subject. The eye movement-related signals were similar in the two ears when saccade direction is defined with respect to the recorded ear. The remaining individual subjects’ data can be found in Supplementary Figure S2.

### Relating the EMREO signal to eardrum displacement

At the low frequencies characteristic of EMREO oscillations (~30 Hz), mean eardrum displacement is directly proportional to the pressure produced in the small volume enclosed between the microphone and the eardrum (~1 cc for humans). We converted the EMREO voltage signal to pressure using the microphone calibration curve. At frequencies below about 200 Hz, the nominal microphone sensitivity quoted by the manufacturer does not apply, and so we used the complex frequency response measured by Christensen et al. (2015; Figure 3; see Methods for details). We found that the group average EMREO had peak-to-peak pressure changes near ~42 mPa for 18 degree contralateral saccades as well as an initial phase opposite to the microphone voltage (Figure 5C, maximum excursion of dark red traces). Using ear-canal dimensions typical of the adult ear (29), this pressure, the equivalent of about 57 dB peak equivalent SPL, corresponds to a mean, peak-to-peak eardrum displacement of ~4 nm. Thus, the inferred initial deflection of the eardrum was always opposite to the direction of the saccade: when the eyes moved left, the eardrums moved right, and vice versa.

**Figure 5.**
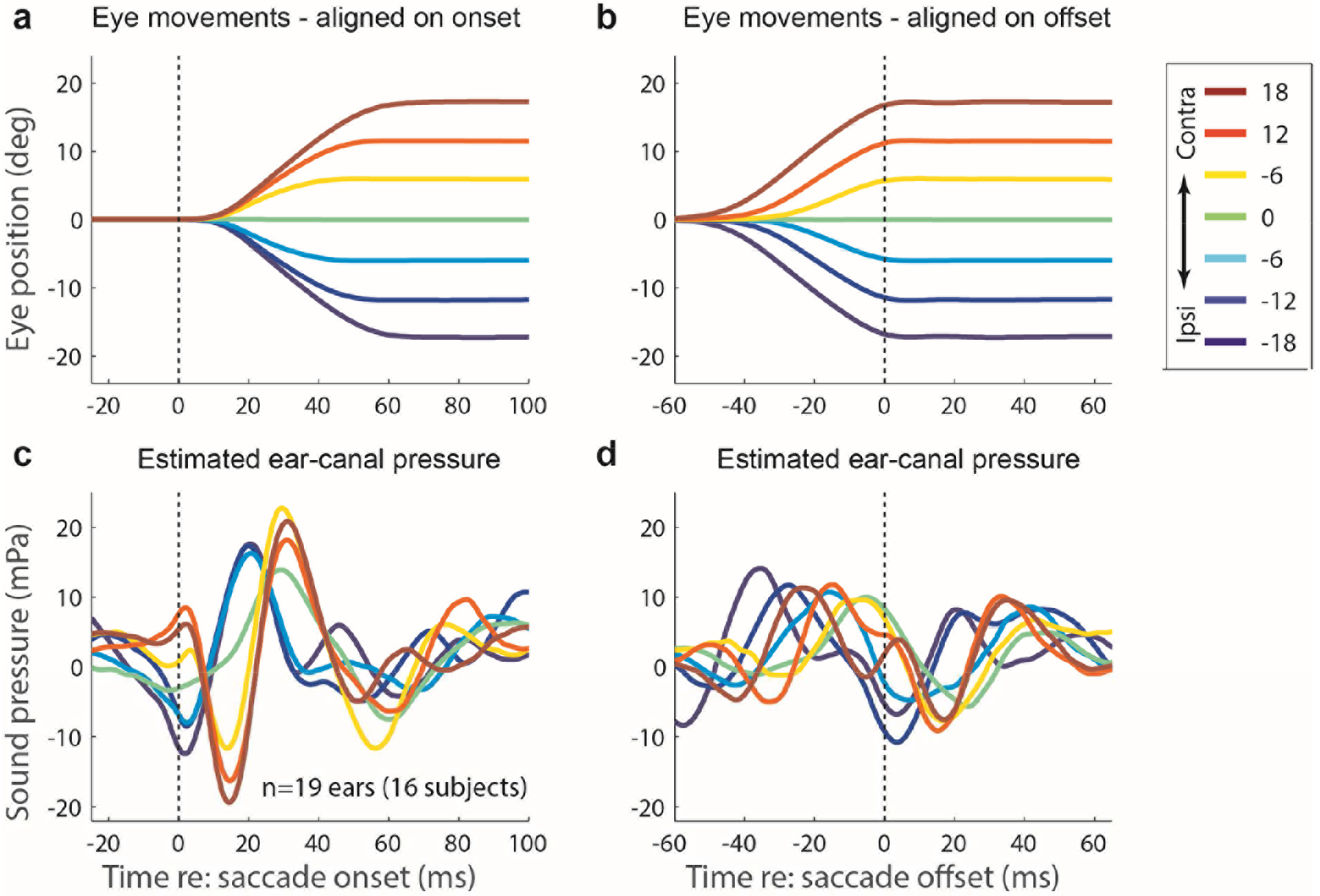
Estimated EMREO pressure for human subjects at saccade onset (a-b) and saccade offset (c-d). Pressures were obtained from the measured microphone voltage using the microphone’s complex frequency response measured at low frequencies by (30).

### EMREOs in monkeys

As noted earlier, eye movements are known to affect neural processing in several areas of the auditory pathway (2, 3, 6, 8, 31–34). Because this previous work was conducted using non-human primate subjects, we sought to determine whether EMREOs also occur in monkeys. We tested the ears of rhesus monkeys (*Macaca mulatta*; *n*=5 ears in 3 monkeys) performing a paradigm involving a trial type that was a hybrid of those used for the humans. The hybrid trial type was silent until after the eye movement, after which a single click was delivered with a 200-270 ms variable delay. EMREOs were observed with timing and waveforms roughly similar to those observed in the human subjects, both at the population level (Figure 6B-D) and at the individual subject level (Figure 6E; Supplemental Figure S3 shows the individual traces). The time-wise regression analysis suggests that the monkey EMREO begins at about the time of the saccade and reaches a robust level about 8 ms later (Figure 4B-E).

**Figure 6.**
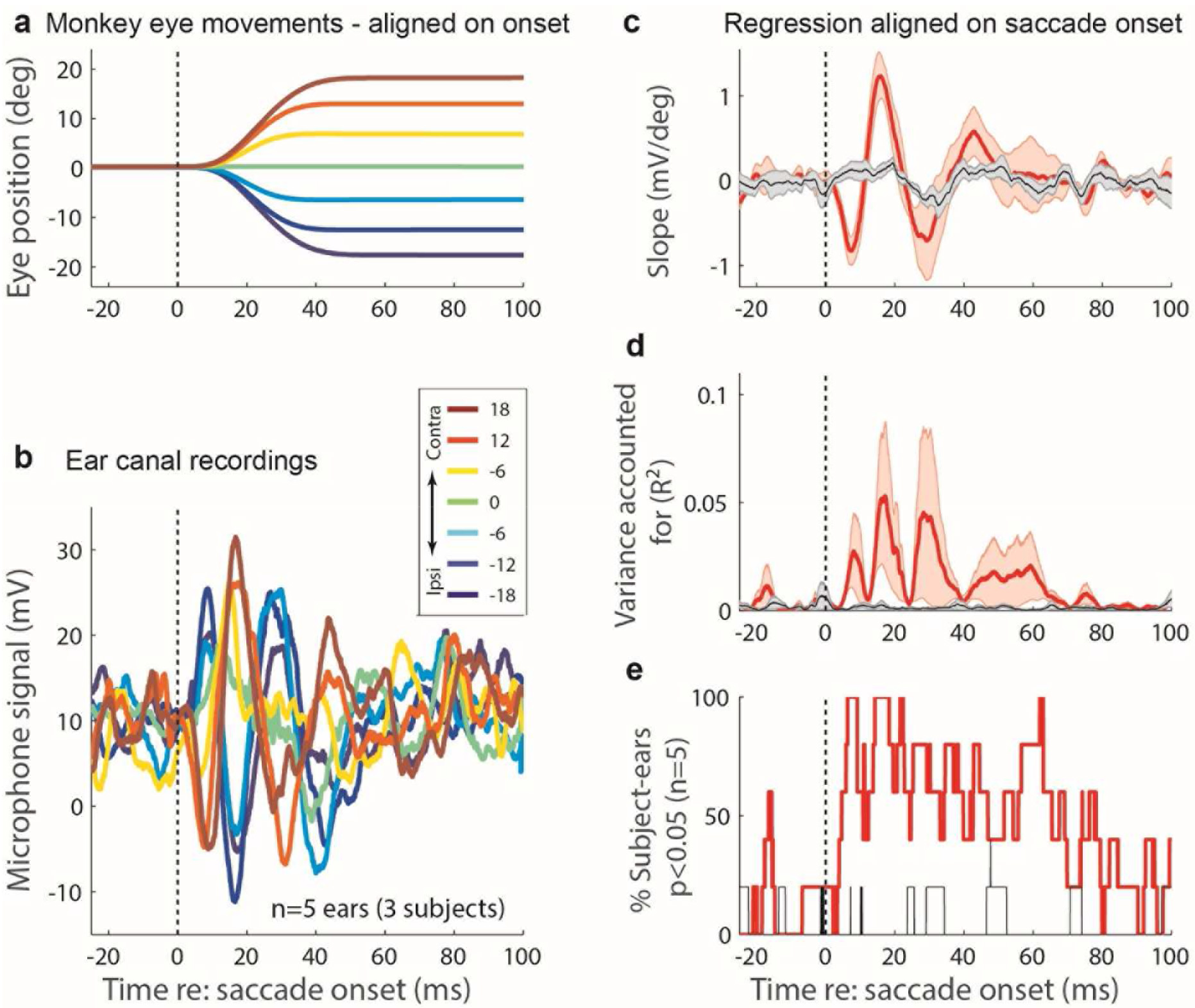
Eye position (**a**), microphone signal of ear-canal pressure (**b**), and results of point-by-point regression (**c-e**) for monkey subject (n=5 ears); all analyses calculated the same way as for the human data (Figures 2-3).

### Controls for electrical artifact

Control experiments ruled out electrical artifacts as possible sources of these observations. In particular, we were concerned that the microphone’s circuitry could have acted as an antenna and been influenced by eye-movement related electrical signals in either the eye-movement measurement system in monkeys (scleral eye coils) and/or some other component of the experimental environment: electrooculographic (EOG) signals resulting from the electrical dipole of the eye, or myogenic artifacts such as electromyographic (EMG) signals originating from the extraocular, facial, or auricular muscles. If such artifacts contributed to our measurements, they should have continued to do so when the microphone was acoustically plugged without additional electrical shielding. Accordingly, we selected four subjects with prominent effects of eye movements on the microphone signal (Figure 7A) and repeated the test while the acoustic port of the microphone was plugged (Figure 7B). The eye movement-related effects were no longer evident when the microphone was acoustically occluded (Figure 7B). This shows that eye movement-related changes in the microphone signal stem from its capacity to measure acoustic signals rather than electrical artifacts. Additionally, EMREOs were not observed when the microphone was placed in a 1-mL syringe (the approximate volume of the ear canal) and positioned directly behind the pinna of one human subject (Figure 7C). In this configuration, any electrical contamination should be similar to that in the regular experiment. We saw none, supporting the interpretation that the regular ear-canal microphone measurements are detecting eardrum motion and not electrical artifact.

**Figure 7.**
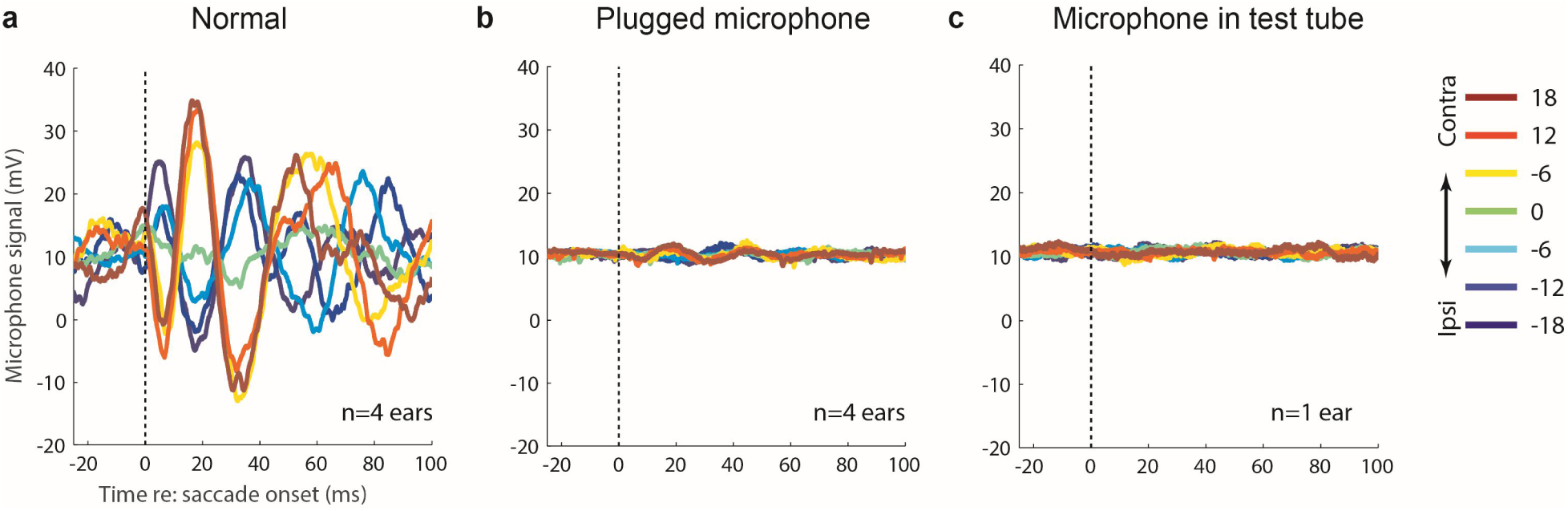
EMREOs recorded normally (**a**) are not observed when the microphone input port is plugged to eliminate acoustic but not electrical contributions to the microphone signal (**b**). The plugged microphone sessions were run as normal sessions except that after calibration, the microphone was placed in a closed earbud before continuing with the session (n=4 subjects). Similarly, EMREOs were not evident when the microphone was placed in a test tube behind a human subject’s head while he/she performed the experiment as normal (**c**).

### EMREOs and click stimuli

EMREOs occurred not only in silence, but also when sounds were presented. During half of our trials for human subjects, acoustic clicks (65 dB peak equivalent SPL) were delivered during the initial fixation, during the saccade to the target, and at both 100 ms and 200 ms after obtaining target fixation. Clicks presented during the saccade superimposed on the ongoing EMREO (Figure 8B1). Subtraction of the clicks measured during the fixation period (Figure 8A1) indicated that the EMREO was not appreciably changed by the presence of the click (Figure 8B2). Unlike saccade onset or offset, the click did not appear to reset the phase of the EMREO.

Clicks delivered after the eye movement was complete (Figure 8C1-D1, 8C2-D2), revealed no new sound-triggered effects attributable to the different static eye fixation positions achieved by the subjects by that point of the trial. Results in monkeys, for which only one post-saccadic click was delivered (at 75 dB peak equivalent SPL), were similar (Figure 8E).

Finally, we tested the peak-to-peak amplitude of the microphone signal corresponding to the click itself. Any changes in the recorded amplitude of a click despite the same voltage having been applied to the earbud speaker would indicate changes to the acoustic impedance of the ear as a function of saccade direction. We calculated the peak-to-peak amplitude of the click as a function of saccade target location, but found no clear evidence of any saccade target location-related differences during any of the temporal epochs (Figure 8A3-E3). If the mechanisms causing the eardrum to move produce concomitant changes in the dynamic stiffness of the eardrum, they are too small to be detected with this technique.

**Figure 8.**
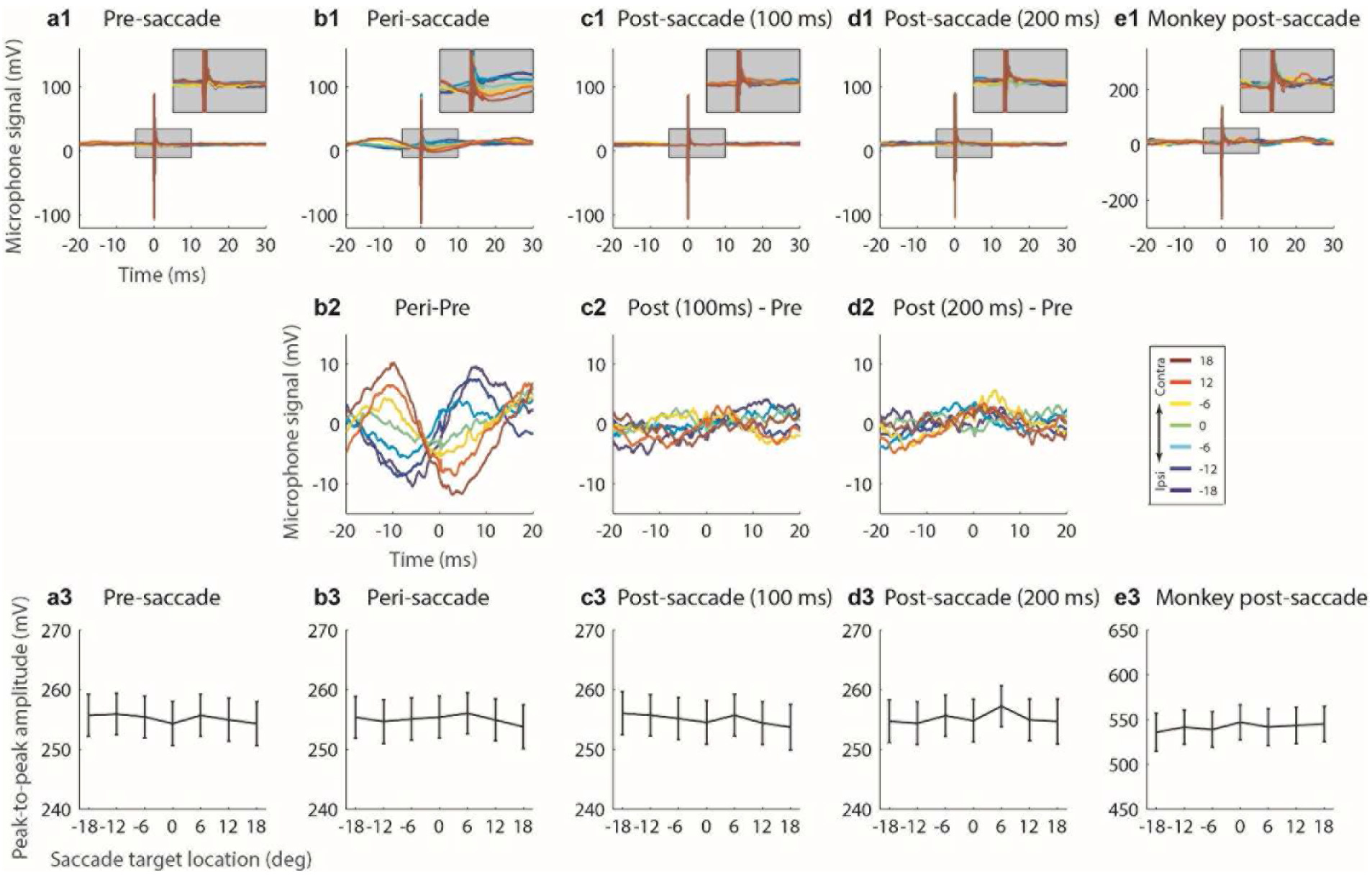
Clicks do not appear to alter EMREOs. Panels a-e, row 1: Mean microphone signals for human subjects (A-D) and monkeys (E) for click presentations at various time points: during the initial fixation (a), during the saccade (approximately 20 ms after saccade onset, b), and 100 ms (c) and 200 ms (d) after target fixation was obtained in humans, and 200-270 ms after target fixation in monkeys (e). The insets zoom in on the peri-click timing for a more detailed view (gray backgrounds). Panels b-d, row 2: residual microphone signal after the first click in each trial was subtracted (human subjects). There were no obvious distortions in the EMREOs at the time of the (removed) click, suggesting that the effects of EMREOs interact linearly with incoming sounds. Panels a-e, row 3: Mean peak-to-peak amplitude of the clicks (Mean ± SE). There were no clear differences in the peak click amplitudes for any epochs, indicating that the acoustic impedance did not change as a function of saccade target location.

## DISCUSSION

Here, we demonstrated a regular and predictable pattern of oscillatory movements of the eardrum associated with movements of the eyes. When the eyes move left, both eardrums initially move right, then oscillate for 3 to 4 cycles with another 1 to 2 cycles occurring after the eye movement is complete. Eye movements in the opposite direction produce an oscillatory pattern in the opposite direction. This eye movement related eardrum oscillation (EMREO) is present in both human and non-human primates and appears similar across both ears within individual subjects.

The impact of the eye movement-related process revealed here on subsequent auditory processing remains to be determined. In the case of otoacoustic emissions (OAEs,), the emissions themselves are easily overlooked, near-threshold sounds in the ear canal. But the cochlear mechanisms that give rise to them have a profound impact on the sensitivity and dynamic range of hearing (35, 36). OAEs thus serve as a biomarker for the health of this important mechanism. EMREOs may constitute a similar biomarker indicating that information about the direction, amplitude, and time of occurrence of saccadic eye movements has reached the periphery and that such information is affecting underlying mechanical processes that may be not themselves be readily observed. EMREO assessment may therefore have value in understanding auditory and visual-auditory deficits that contribute to disorders ranging from language impairments to autism and schizophrenia as well as multisensory deficits associated with traumatic brain injuries.

One important unresolved question is whether the oscillatory nature of the effect observed here reflects the most important aspects of the phenomenon. It may be that the oscillations reflect changes to internal structures within the ear that persist statically until the eye moves again. If so, then the impact on auditory processing would not be limited to the period of time when the eardrum is actually oscillating. Instead, processing could be affected at all times, updating to a new state with each eye movement.

If the oscillations themselves are the most critical element of the phenomenon, and auditory processing is only affected during these oscillations, then it becomes important to determine for what proportion of the time they occur. In our paradigm, we were able to demonstrate oscillations occurring over a total period of ~110 ms, based on the aggregate results from aligning on saccade onset and offset. They may continue longer than that, but with a phase that changes in relation to some unmeasured or uncontrolled variable. As time passes after the eye movement, the phase on individual trials might drift such that the oscillation disappears from the across-trial average. But with the eyes moving about three times per second under normal conditions, an EMREO duration of about 110 ms for each saccade corresponds to about 1/3 of the time in total.

Regardless, evidence arguing for a possible influence on auditory processing can be seen in the size of EMREOs. A head-to-head comparison of EMREOs with OAEs is difficult since auditory detection thresholds are higher in the frequency range of EMREOs than in the range of OAEs, but EMREOs appear to be at least comparable and possibly larger than OAEs. For 18 degree horizontal saccades, EMREOs produce a maximum peak-equivalent pressure of about 57 dB SPL, whereas click-evoked otoacoustic emissions (OAEs) range from ~0-20 dB SPL in healthy ears. The similar or greater scale of EMREOs supports the interpretation that they too likely reflect underlying mechanisms of sufficient strength to make a meaningful contribution to auditory processing.

In whatever fashion EMREOs or their underlying mechanism contribute to auditory processing, we surmise that the effect concerns the localization of sounds with respect to the visual scene or using eye movements. Determining whether a sight and a sound arise from a common spatial position requires knowing the relationship between the visual eye-centered and auditory head-centered reference frames. Eye position acts as a conversion factor from eye-to head-centered coordinates (1), and eye movement related signals have been identified in multiple auditory-responsive brain regions (2–17). The direction/phase and amplitude of EMREOs contain information about the direction and amplitude of the accompanying eye movements, which situates them well for playing a causal role in this process.

Note that we view the EMREO as most likely contributing to *normal and accurate* sound localization behavior. Sounds can be accurately localized using eye movements regardless of initial eye position (37), and this is true even for brief sounds presented when the eyes are in flight (38). Although some studies have found subtle influences of eye position on sound perception tasks (39–46), such effects are only consistent when prolonged fixation is involved (47–49). Holding the eyes steady for a seconds to minutes represents a substantial departure from the time frame in which eye movements normally occur. It is possible that deficiencies in the EMREO system under such circumstances contribute to these inaccuracies.

The source of the signals that cause EMREOs is not presently known. Because the EMREOs precede, or occur simultaneously with actual movement onset, it appears likely that they derive from a copy of the motor command to generate the eye movement, rather than a proprioceptive signal from the orbits, which would necessarily lag the actual eye movement. This centrally-generated signal must then affect the activity of either MEMs, OHCs, or a combination of both.

Based on similarities between our recordings and known physiological activity of the MEMs, we presume that the mechanism behind this phenomenon is most likely the MEMs. Specifically, the frequency (around 20-40 Hz) is similar to oscillations observed in previous recordings of both the tensor tympani and stapedius muscles (50–55). Furthermore, it seems unlikely, given the measured fall-off in reverse middle-ear transmission at low frequencies (56), that OHC activity could produce ear-canal sounds of the magnitude observed (42 mPa peak-to-peak, or 57 dB peak-equivalent SPL). Although MEMs are often associated with bilaterally attenuating loud environmental and self-generated sounds (which their activity may precede), they are also known to be active in the absence of explicit auditory stimuli, particularly during rapid eye movement (REM) sleep (57–60),non-acoustic startle reflexes (55, 61–65), and have been found to exhibit activity associated with movements of the head and neck in awake cats (61, 66).

More generally, we have demonstrated that multisensory interactions occur at the most peripheral possible point in the auditory system and that this interaction is both systematic and substantial. This observation builds on studies showing that attention—either auditory-guided or visually-guided—can also modulate the auditory periphery (21–27). Our findings also raise the intriguing possibility that efferent pathways in other sensory systems—for instance, those leading to the retina (67–77)—also carry multisensory information to help refine peripheral processing. This study contributes to an emerging body of evidence suggesting that the brain is best viewed as a dynamic system in which top-down signals modulate feed-forward signals. That is, the brain integrates top down information early in sensory processing to make the best informed decision about the world with which it has to interact.

## METHODS

### Human Subjects and Experimental Paradigm

#### Human data set I

Human subjects (*n*=16, 8 females, aged 18-45 years; participants included university students as well as young adults from the local community) were involved in this study. All procedures involving human subjects were approved by the Duke University Institutional Review Board. Subjects had apparently normal hearing and normal or corrected vision. Informed consent was obtained prior to testing, and all subjects received monetary compensation for participation. Stimulus (visual and auditory) presentation, data collection, and offline analysis were run on custom software utilizing multiple interfaces (behavioral interface and visual stimulus presentation: Beethoven software [eye position sampling rate: 500Hz], Ryklin Inc.; auditory stimulus presentation and data acquisition: Tucker Davis Technologies [microphone sampling rate: 24.441 kHz]; data storage and analysis: Matlab, Mathworks).

Subjects were seated in a dark, sound attenuating room. Head movements were minimized using a chin rest, and eye movements were tracked with an infrared camera (EyeLink 1000 Plus). Subjects performed a simple saccade task (Figure 2A,B). The subject initiated each trial by obtaining fixation on an LED located at 0° in azimuth and elevation and about 2 meters away. After 200 ms of fixation, the central LED was extinguished and a target LED located 6° above the horizontal meridian and ranging from -24° to 24°, in 6° intervals, was illuminated. The most eccentric targets (±24°) were included despite being slightly beyond the range at which head movements normally accompany eye movements, which is typically 20° (78, 79), because we found in preliminary testing that including these locations improved performance for the next most eccentric locations, i.e. the ±18 ° targets. However, because these (±24°) locations were difficult for subjects to perform well, we presented them on only 4.5% of the trials (in comparison to 13% for the other locations), and we excluded them from analysis. After subjects made saccades to the target LED, they maintained fixation on it (9° window diameter) for 250 ms until the end of the trial. If fixation was dropped (i.e., if the eyes traveled outside of the 9° window at any point throughout the initial or target fixation periods), the trial was terminated and the next trial began.

On half of the trials, task-irrelevant sounds were presented via the earphones of an earphone/microphone assembly (Etymotic 10B+ microphone with ER 1 headphone driver) placed in the ear canal and held in position through a custom molded silicone earplug (Radians Inc.). These sounds were acoustic clicks at 65 dB peak-equivalent SPL, produced by brief electrical pulses (40 μs positive monophasic pulse) and were presented at four time points within each trial: during the initial fixation period (100 ms after obtaining fixation); during the saccade (approximately 20 ms after initiating an eye movement); 100 ms after obtaining fixation on the target; and 200 ms after obtaining fixation on the target.

Acoustic signals from the ear canal were recorded via the in-ear microphone throughout all trials, and were recorded from one ear in 13 subjects (left/right counterbalanced) and from both ears in separate sessions in the other 3 subjects, for a total of *n*=19 ears tested. Testing for each subject ear was conducted in two sessions over two consecutive days or within the same day but separated by a break of at least one hour in between sessions. Each session involved about 600 trials and lasted a total of about 30 minutes. The sound delivery and microphone system was calibrated at the beginning of every session using a custom script (Matlab) and again after every block of 200 trials. The calibration routine played a nominal 80 dB SPL sound—a click, a broadband burst, and a series of tones at frequencies ranging from 1 to 12 kHz in 22 steps—in the ear canal and recorded the resultant sound pressure. It then calculated the difference between the requested and measured sound pressures and calculated a gain adjustment profile for all sounds tested. Auditory recording levels were set with a custom software calibration routine (Matlab) at the beginning of each data collection block (200 trials). All conditions were randomly interleaved.

#### Human Data Set II

Additional data were collected from 8 ears of 4 new subjects (ages 20-27, 3 female, 1 male, 1 session per subject) to verify that the relationship between microphone voltage and saccade amplitude/target location held up when sampled more finely within a hemifield. All procedures were the same as in the first data set with three exceptions: 1) the horizontal target positions ranged from -20° to +20° degrees in 4° increments (the vertical component was still 6°); 2) the stimuli were presented from a color LCD monitor (70cm ◻ 49cm) at a distance of 85cm; and 3) no clicks were played during trials. The microphone signal was also calibrated differently: clicks played between trials revealed the stability of its impulse response and overall gain during and across experimental sessions. This helped identify decreasing battery power to the amplifiers, occlusion of the microphone barrel, or a change in microphone position. A Focusrite Scarlett 2i2 audio interface was used for auditory stimulus presentation and data acquisition [microphone sampling rate 48kHz]. Stimulus presentation and data acquisition were controlled through custom Matlab scripts using The Psychophysics Toolbox extension (80–82) and the Eyelink Matlab toolbox (83).

These data are presented in Supplementary Figure 1; the remainder of the human figures derive from data set I.

### Monkey Subjects and Experimental paradigm

All procedures conformed to the guidelines of the National Institutes of Health (NIH Pub. No. 86-23, Revised 1985) and were approved by the Institutional Animal Care and Use Committee of Duke University. Monkey subjects (*n*=3, all female) underwent aseptic surgical procedures under general anesthesia to implant a head post holder to restrain the head and a scleral search coil (Riverbend Eye Tracking System) to track eye movements (80, 81). After recovery with suitable analgesics and veterinary care, monkeys were trained in the saccade task described above for the human subjects. Two monkeys were tested with both ears in separate sessions, whereas the third monkey was tested with one ear, for a total of *n*=5 ears tested.

The trial structure was similar to that used in human data set I but with the following differences (Figure 2B, red traces): (1) eye tracking was done with a scleral eye coil; (2) task-irrelevant sounds were presented on all trials, but only one click was presented at 200-270 ms after the saccade to the visual target (the exact time jittered in this range, and was therefore at the same time or slightly later than the timing of the fourth click for human subjects); (3) the ±24° targets were presented in equal proportion to the other target locations, but were similarly excluded from analysis as above; in addition there were targets at ±9° which were also discarded as they were not used with the human subjects; (4) initial fixation was 110-160 ms (jittered time range), while target fixation duration was 310-430 ms; (5) monkeys received a fluid reward for correct trial performance; (6) disposable plastic ear buds containing the earphone/microphone assembly as above were placed in the ear canal for each session; (7) auditory recording levels were set at the beginning of each data collection session (the ear bud was not removed during a session, and therefore no recalibration occurred within a session); (8) sessions were not divided into blocks (the monkeys typically performed consistently throughout an entire session and dividing it into blocks was unnecessary); and (9) the number of trials per session was different for monkeys versus humans (which were always presented exactly 1200 trials over the course of the entire study); the actual number of trials performed varied based on monkey’s performance tolerance and capabilities for the day. Monkeys MNN012 and MYY002 were both run for 4 sessions per ear (no more than one session per day) over the course of 2 weeks; MNN012 correctly performed an average of 889 out of 957 trials at included target locations per session for both ears, while MYY002 correctly performed 582 out of 918 trials per session on average. Monkey MHH003 was only recorded one day and yielded 132 correct out of 463 trials. Despite this much lower performance, visual inspection of the data suggested they were consistent with other subject data and were therefore included for analysis. It is worth highlighting that the effect reported in this paper can be seen, in this case, with comparatively few trials.

### Control sessions

To verify that the apparent effects of eye movements on ear-generated sounds were genuinely acoustic in nature and did not reflect electrical contamination from sources such as myogenic potentials of the extraocular muscles or the changing orientation of the electrical dipole of the eye ball, we ran a series of additional control studies. Some subjects (*n*=4 for plugged microphone control; *n*=1 for syringe control) were invited back as participants in these control studies.

In the first control experiment, the microphone was placed in the ear canal and subjects performed the task but the microphone’s input port was physically plugged, preventing it from detecting acoustic signals (see Figure 7). This was accomplished by placing the microphone in a custom ear mold in which the canal-side opening for the microphone was blocked. Thus, the microphone was in the same physical position during these sessions, and should therefore have continued to be affected by any electrical artifacts that might be present, but its acoustic input was greatly attenuated by the plug. Four subject ears were re-tested in this paradigm in a separate pair of data collection sessions from their initial “normal” sessions. The control sessions were handled exactly as the normal sessions except that the plugged ear mold was used to replace the regular open ear mold after the microphone was calibrated in the open ear. Calibration was executed exactly as in the normal sessions prior to plugging the ear mold.

In the second control, trials were run exactly as a normal session except that the microphone was set into a 1 mL syringe [approximately the average volume of the human ear canal (82)] using a plastic ear bud, and the syringe was place on top of the subject’s ear behind the pinna. Acoustic recordings were taken from within the syringe while a human subject executed the behavioral paradigm exactly as normal. Sound levels were calibrated to the syringe at the start of each block.

### Data Analysis

#### Initial Processing

Unless specifically stated otherwise, all data are reported using the raw voltage recorded from the Etymotic microphone system. Results for individual human and monkey subjects were based on all of that subject’s correct and included trials (with ±24° and 9 ° target locations excluded, the average number of correct trials per human subject-ear for each session was 150, for a total of 900 across the 6 sessions).

#### Trial exclusion criteria

Trial exclusion criteria were based on saccade performance and microphone recording quality. For saccade performance, trials were excluded if: a) the reaction time to initiate the saccade was >250 ms; or b) the eyes deviated from the path between the fixation point and the target by more than 9°. These criteria resulted in the exclusion of 18.5% ±11.1% per ear recorded.

Microphone recording quality was examined for click and no-click trials separately, but followed the same guidelines. The mean and standard deviation of microphone recordings over a whole block were calculated from all successful trials (after exclusions based on saccade performance). If the standard deviation of the voltage values of a given trial was more than 3 times the standard deviation of the voltage values across the whole block, it was excluded. Sample-wise z-scores were calculated relative to the mean and standard deviation of successful trials within a block. Trials with 50 samples or more with z-scores in excess of 10 were also excluded. These requirements removed trials containing noise contamination from bodily movements (like swallowing and heavy breaths) or from external noise sources (acoustic and electric). These two criteria excluded an average of 1.7% ± 1.5% of trials that had passed the saccade exclusion criteria. Overall, 20% of trials were excluded because of saccade performance or recording quality.

#### Saccade-microphone synchronization

Eye-position data were median filtered over three samples (4 ms), resampled to the microphone sampling rate (from 500 Hz to 24.5 kHz), and smoothed with a moving average filter over a 1 ms window. This process minimized the compounding error of non-monotonic recording artifacts after each successive differentiation of eye-position to calculate eye-velocity, acceleration, and jerk (the time-derivative of acceleration).

Microphone data were synchronized to two time points in the saccades: initiation and termination. Saccade onsets were determined by locating the time of the first peak in the jerk of the saccades. These onset times for each trial were then used to synchronize the microphone recordings at the beginning of saccades. Because saccades of different lengths took different amounts of time to complete and because performance of saccades to identical targets varied between trials, saccade completions occurred over a range of tens of milliseconds. As a result, recordings of eardrum behavior related to the later stages of saccades would be poorly aligned if synchronized by saccade onset alone. Therefore, saccade termination, or offset, was determined by locating the second peak in the saccade jerk and used to synchronize microphone recordings with saccade completion.

#### Statistical analyses

We evaluated the statistical significance of the effects of eye movements on signals recorded with the ear canal microphone with a regression analysis at each time point (microphone signal vs. saccade target location), for each subject. This produced a time series of statistical results (slope, *R*^2^, and *p*-value; see Figure 2E-G). This analysis was performed twice: first with all trials synchronized to saccade onset and second with all trials synchronized to saccade offset.

Given that the dependence of one sample relative to the next sample was unknown, a post-hoc correction (e.g., Bonferroni correction) was not practical. Instead, we used a Monte Carlo technique to estimate the chance-related effect size and false positive rates of this test. We first scrambled the relationship between each trial and its saccade target location assignment, then ran the same analysis as before. This provided an estimate of how our results should look if there was no relationship between eye movements and the acoustic recordings.

Peak click amplitudes were calculated for each trial involving sounds by isolating the maximum and minimum peaks during the click stimuli. This allowed us to look for possible changes in acoustic impedance in the ear canal.

#### Estimation of Eardrum Motion

The sensitivity of the microphone used in the experiments (Etymotic Research ER10B+) rolls off at frequencies below about 200 Hz, where it also introduces a frequency-dependent phase shift (30). Consequently, the microphone voltage waveform is not simply proportional to the ear-canal pressure (or eardrum displacement) at frequencies characteristic of the EMREO waveform. To obtain the ear-canal pressure, and estimate the eardrum motion that produced it, we converted the microphone voltage to pressure using published measurements of the microphone’s complex-valued frequency response (30).

Because the measured frequency response (*H*_mic_, with units of V/Pa) was sampled at 48 kHz, we first transformed it into the time-domain representation of the microphone’s impulse response, resampled the result at our sampling rate of 24.4 kHz, and then re-transformed it back into the frequency domain as *H*_mic′_. For each trial, the Fast Fourier transform of microphone voltage was divided by *H*_mic′_ and the inverse Fast Fourier transform was then calculated to produce the estimate of ear-canal, *P*_ec_.

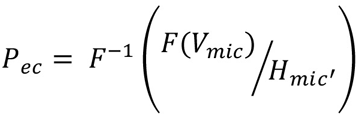

Eardrum displacement *x*_ed_ was computed from the measured pressure using the equation *x*_ed_ = (*V*/*A*) *P*_ec_/*ρ*_0_*c*^2^, where *V* is the volume of the residual ear-canal space (~2 cc), *A* is the cross-sectional area of the eardrum (~60 mm^2^), *ρ*_0_ is the density of air, and *c* is the speed of sound.

## SUPPLEMENTARY FIGURES

**Supplementary Figure 1.**
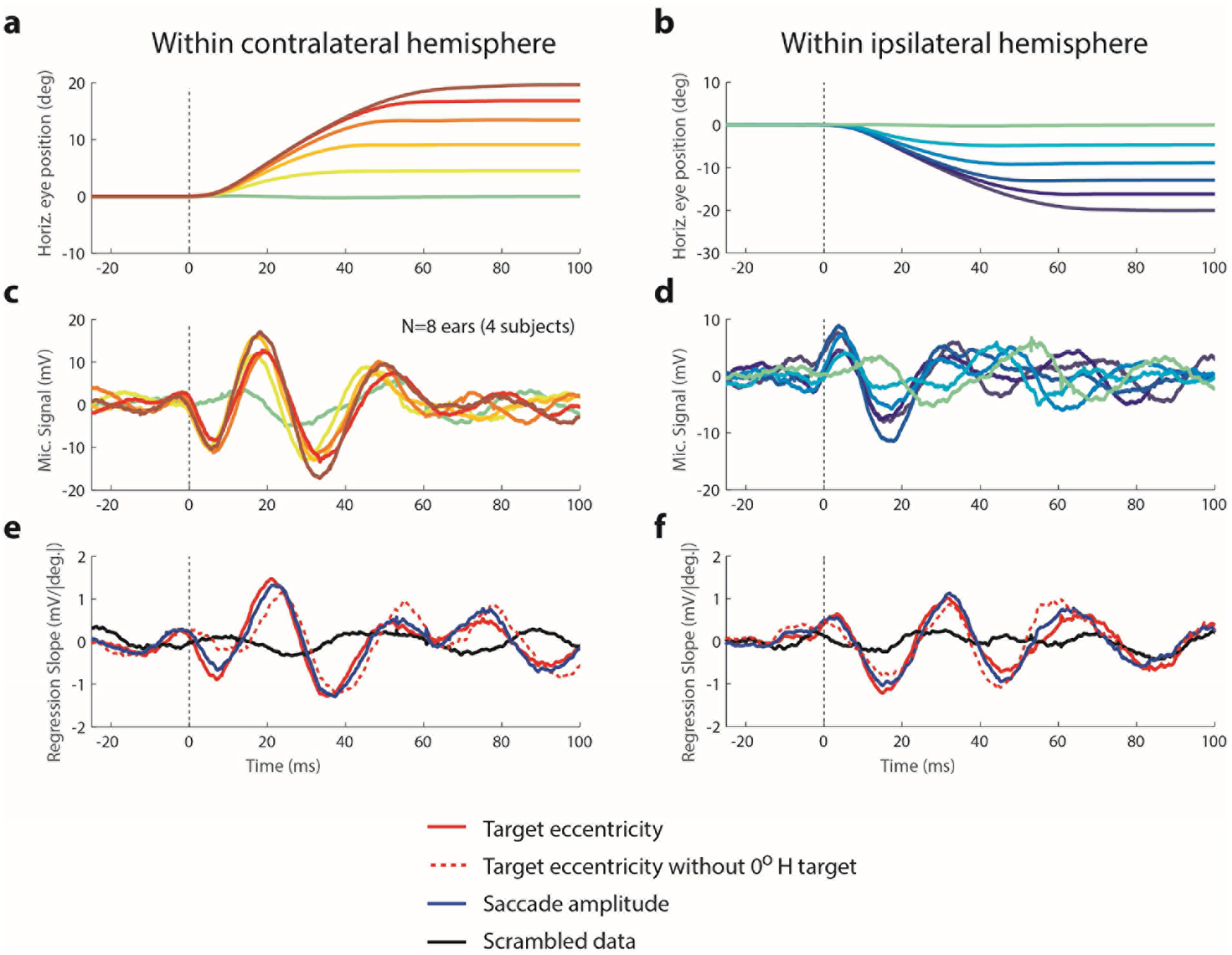
We repeated the basic experiment in an additional 8 ears in 4 subjects, sampling target locations more finely and analyzing the data within the contralateral (A, C, E) or within the ipsilateral hemifields (B, D, F). Panels A, B show the average horizontal component of saccades to the targets at locations 0, 4, 8, 12, 16, and 20 degree targets in the contralateral (A) or ipsilateral (B) fields (aligned on saccade onset as in Figure 2C; all saccades had a vertical component of 6 degrees as well as their horizontal component). Panels C, D show the average microphone voltages associated with these saccades (as in Figure 2E), and panels E, F show the slope of the regression relating the microphone voltage in each sample to the target eccentricity (red traces) or saccade amplitude (blue trace) on that trial in comparison to one scrambled run (red vs. black traces). The dashed red trace illustrates the slope if the results involving the target with only a vertical component (0 deg horizontal, green traces in A-B, D-E) are omitted from the regression. All three analysis variations produced similar results.

**Supplementary Figure 2.**
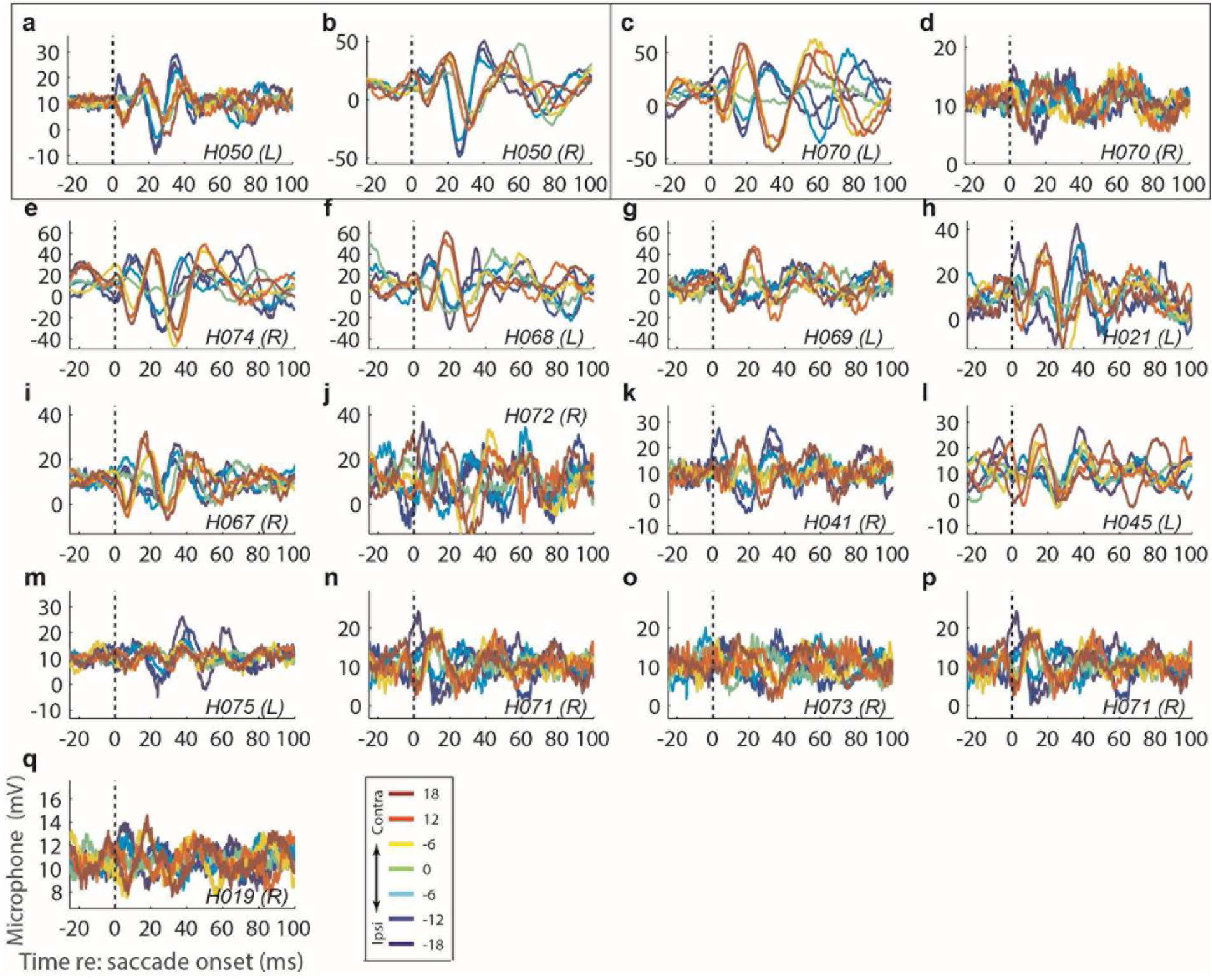
Data from individual human subject ears. Conventions the same as Figure 4. Panels a-b and c-d, enclosed in boxes, represent data from different ears in the same two subjects. Exploratory analysis in a follow-up study suggests variation in microphone placement within the ear canal may contribute to apparent individual differences in EMREO signal-to-noise ratio.

**Supplementary Figure 3:**
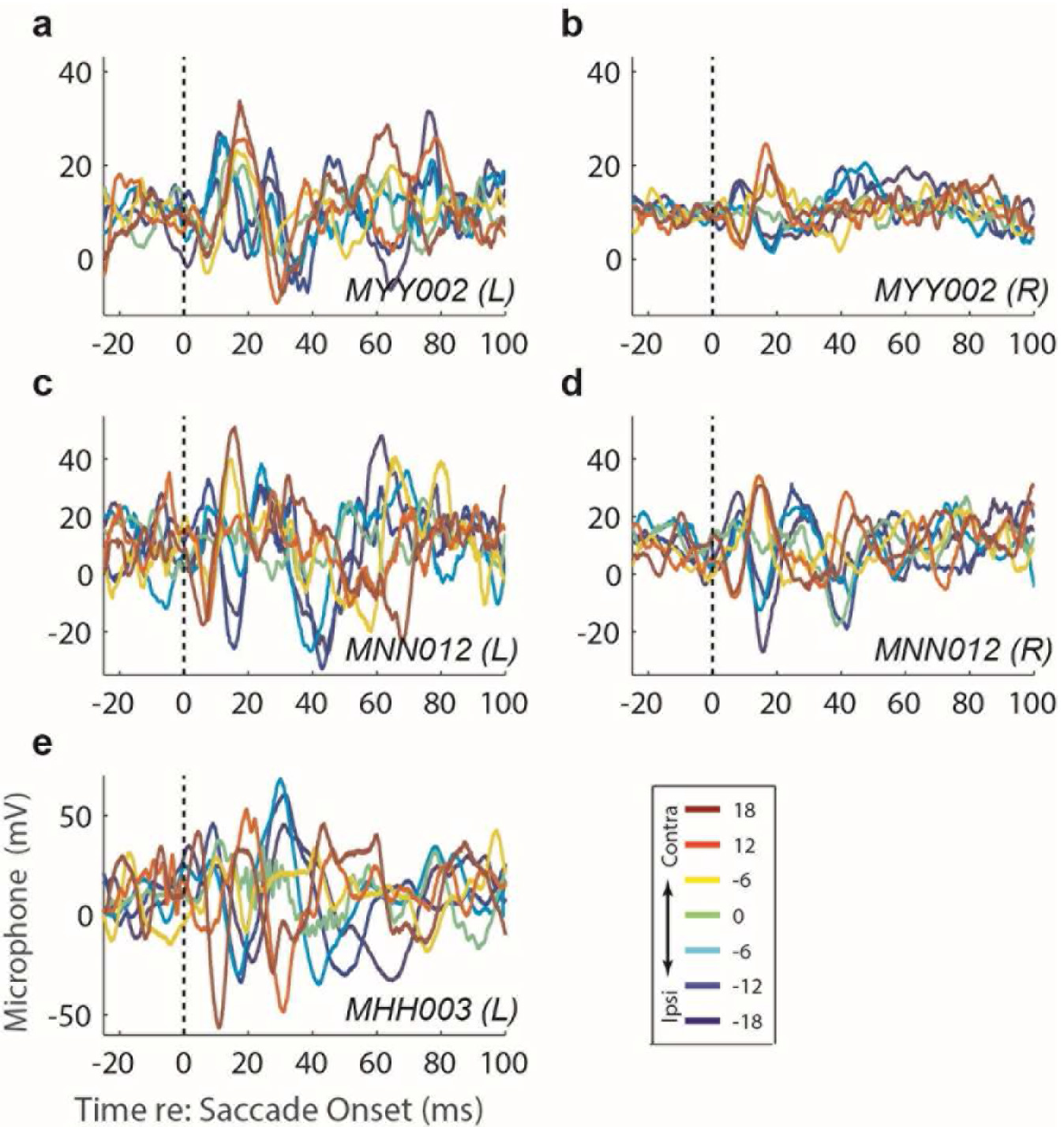
Individual subject results for all monkey ears. Analysis details are the same as individual human subjects (Supplementary Figure S2).

## Acknowledgments

We are grateful to Tom Heil, Jessi Cruger, Karen Waterstradt, Christie Holmes, and Stephanie Schlebusch for technical assistance. We thank Marty Woldorff, Jeff Beck, Tobias Overath, Barbara Shinn-Cunningham, Valeria Caruso, Daniel Pages, Shawn Willett, Jeff Mohl for numerous helpful discussions and comments during the course of this project.

